# Development and validation of an fMRI-informed EEG model of reward-related ventral striatum activation

**DOI:** 10.1101/2022.11.01.514407

**Authors:** Neomi Singer, Gilad Poker, Netta Dunsky, Shlomi Nemni, Maayan Doron, Travis Baker, Alain Dagher, Robert J Zatorre, Talma Hendler

## Abstract

Reward processing is essential for our mental-health and well-being. Here, we present the development and validation of a scalable fMRI-informed EEG model related to reward processing in the ventral-striatum (VS); a central reward circuit node. Simultaneous EEG/fMRI data were acquired from 17 healthy individuals listening to pleasurable music, and used to construct a one-class regression model for predicting the reward-related VS-BOLD signal using spectro-temporal features from the EEG. Validation analyses, applied on EEG/fMRI data from a different group (N=14), revealed that the EEG model predicted VS-BOLD activation from the simultaneous EEG to a greater extent than a model derived from another anatomical region. The VS-EEG-model was also modulated by musical pleasure and predictive of the VS-BOLD during a monetary reward task, further indicating it functional relevance. These findings provide compelling evidence for the use of a scalable yet precise EEG-only probe of VS-originated reward processing, which could serve for process specific neruo-monitoring and -modulation.

## 1. Introduction

The assignment of **reward** value is a central driving force of human behavior, and is thus critical for mental health and well-being. Since the seminal self-stimulation animal studies carried out over 60 years ago (Olds and Milner, 1954), a large body of pharmacological and physiological studies on animals has pointed to the critical role of brain structures along the mesolimbic pathway, especially the ventral striatum (VS) and the dopaminergic midbrain (Ventral Tegmental area; VTA), in reward processing (Berridge and Robinson, 2003; Cardinal et al., 2002; Haber and Knutson, 2009; Schultz et al., 1997). Balanced functioning of central nodes within this circuitry is considered essential for motivation-related processes, including consumption of desirable objects (‘wanting’), the experience of pleasure obtained from rewards (‘liking’), and learning predictive contingencies that signal reward for future encounters (‘learning’) (Berridge and Robinson, 2003).

Non-invasive neuroimaging techniques, such as functional Magnetic Resonance Imaging (fMRI) and Positron Emission Tomography (PET), have also supported these animal model observations in humans (Bartra et al., 2013; Haber and Knutson, 2009; Jauhar et al., 2021; Knutson and Greer, 2008; Liu et al., 2011). This body of work revealed that major nodes in the ‘reward circuitry,’ such as the VS and ventromedial PreFrontalCortex (vmPFC), are involved in the processing of primary and secondary rewards, ranging from food, sex and monetary gains (Sescousse et al., 2013), to more complex, abstract stimuli, including even music (Blood and Zatorre, 2001; Koelsch, 2020; Mas-Herrero et al., 2021; Menon and Levitin, 2005; Salimpoor et al., 2013; Shany et al., 2019). These studies further highlight local dopamine release as one of its major correlates (Hakyemez et al., 2008; Salimpoor et al., 2011). Importantly, disturbances of this dopaminergic circuit are associated with psychiatric symptoms of hampered reward processing, such as anhedonia (Pizzagalli, 2014), apathy (Kirschner et al., 2020), and addiction (Luijten et al., 2017). Therefore, probing the neural activation within this circuit could be valuable for basic science and clinical application in neuropsychiatry.

The best non-invasive way to probe the reward system, including its deep-brain regions, is with neuroimaging such as fMRI (Rosen and Savoy, 2012). However, this technique is not scalable due to poor accessibility, as it is limited to specific academic and clinical sites and is expensive. Electroencephalography (EEG), on the other hand, is low-cost and accessible, thus optimal for dense monitoring in more natural settings or for clinical applications. However, EEG suffers from a poor spatial resolution that especially hampers targeting deep-brain areas, such as the VS. To overcome this spatial limitation, computational tools can be used. For example, brain source estimation approaches that rely on biophysical assumptions of distributed source modeling, such as standardized or exact Low-Resolution Electromagnetic Tomography (eLORETA; Pascual-Marqui, 2007, respectively, sLORETA; 2002), have been popular techniques for estimating the location of brain areas that generate measured scalp EEG by addressing the inverse problem (Michel and Brunet, 2019). Other models that focus on addressing the forward problem using head models such as the Boundary Element Method (BEM) rely on head anatomy to estimate the projected signals from various regions (Akalin-Acar and Gençer, 2004). However, these various EEG modeling solutions necessitate the use of a dense grid of electrodes and/or precise information about head anatomy via an individual’s structural MRI, limiting the potential generalizability, mobility and scalability of this method (Attal et al., 2012).

Given the complementary nature of EEG and fMRI techniques, they may be integrated for source localization (Abreu et al., 2018). Theory-driven approaches that utilize fMRI to improve EEG localization have attempted to construct a forward model that traces neuronal activity from both measures (Valdes-Sosa et al., 2009). However, such a model-based approach relies on a-priori assumptions regarding the biophysical origins of the common source of EEG and fMRI signals.

To overcome the inherent limitation of lacking a-priori anatomical knowledge, and to avoid the ill-posed inverse problem of detecting activated sources, it is possible, when targeting a specific region, to apply advanced statistical tools to benefit from the integration between the EEG and fMRI by predicting the fMRI activation related to a particular region of interest (ROI) based on the simultaneously acquired EEG signal. Such a statistical modeling-based framework, which capitalizes on the advantages of EEG and fMRI to probe activation in a particular region of interest, has been demonstrated to be effective most recently for the amygdala (*32*). This modeling approach utilizes machine learning to generate a generic, one-class (subject-independent) fMRI-informed EEG model of the activation within a particular region (2016, 2014). This procedure results in an EEG model that is based on weights of different frequency bands and their associated time delays, enabling the prediction of blood-oxygen-level-dependent (BOLD) activation signals in the targeted region using EEG alone (termed Electrical FingerPrint: EFP). This amygdala-BOLD informed model has been validated using simultaneous EEG/fMRI data of the same individuals from different sessions (Meir-Hasson et al., 2016), and of a different group of individuals, revealing it to be a reliable predictor of the BOLD activity in the amygdala and connected network (Keynan et al., 2016). The algorithmic development of such cross-modal prediction inspired the construction of predictive models of additional regions by others, such as the motor cortex (Rudnev et al., 2021), and groups of regions such as the facial expression processing network (Simões et al., 2020).

The fMRI-informed EEG modeling approach may be especially valuable for developing EEG probes that target subcortical brain activity, which is reliably represented by fMRI activity, such as that induced in a reward-related context (Lubianiker et al., 2019). Yet all the previous attempts were driven solely by anatomical consideration of the fMRI activity. The project hereby presented aims to employ this methodology to develop, test, and validate an EEG model based on a process-related BOLD activation in a core reward circuit node – the VS. To ensure that the modeling be performed within a process-specific context of reward (Lubianiker et al., 2019), we acquired simultaneous EEG/fMRI data from 17 participants while they were listening to individually-tailored highly pleasurable music excepts, a task known to modulate the reward circuitry, and particularly the VS (Salimpoor et al., 2011). We used the dataset from this cohort, which is termed the ‘modeling cohort’, to construct a one-class model of VS-BOLD activation from the time-frequency features of the EEG data (hereby termed VS-EFP; Fig. 1). We then turned to assess the performance of the model using the leave-one-out cross-validation approach applied on the modeling dataset, as well as using external validation on an independent dataset from a ‘validation cohort’ – a different group of 14 participants who underwent the same task procedure during fMRI/EEG acquisition. In both cohorts, we estimated the sensitivity and anatomical specificity of the VS-EFP model in predicting the BOLD activity of the VS and additional reward-related regions. We further tested the functional validity of the VS-EFP model by examining whether the EFP signal is modulated by musical reward, and if it is also predictive of VS-BOLD activation under another well-established task that is known to modulate the VS, namely, the Monetary Incentive Delay (MID) task (Knutson et al., 2000; Lutz and Widmer, 2014).

**Fig. 1.**
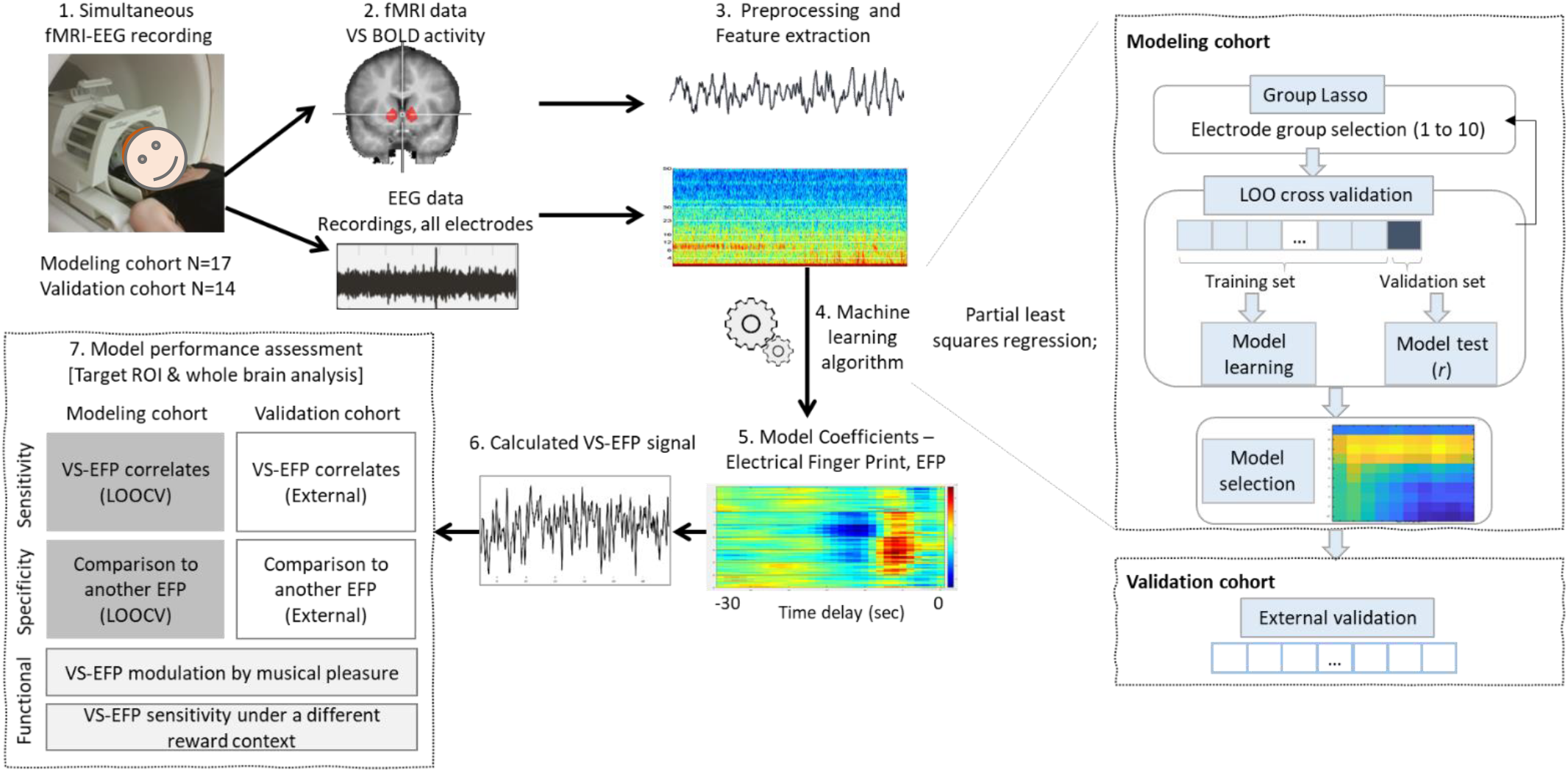
Scheme illustrating the construction of the fMRI-informed EEG model of ventral striatal activity; hereby termed VS-Electrical Finger Print (VS-EFP). 1. EEG and fMRI data were acquired simultaneously during a pleasurable music listening task (19). 2. The fMRI time course from a selected region of interest in the bilateral VS, and 3. time-frequency matrix obtained from the EEG data, were used to calculate the model using a 4. machine learning procedure. 5. The model’s coefficient matrix was applied on the EEG data to construct the 6. VS-EFP time-series. 7. The resulting time series were used to assess the model’s sensitivity, neuroanatomical specificity, and functional validity via ROI and whole-brain random effects analyses. Initial evaluation of the model’s performance in terms of sensitivity and specificity was carried out on the modeling dataset, using the leave on out cross validation approach. Validation of the model was then carried out by examining its performance on the validation cohort; an independent data-set of a completely different group of 14 participants who underwent the same EEG-FMRI scanning procedure. **Abbreviations**: VS=ventral striatum; EFP = Electrical FingerPrint; fMRI = functional Magnetic Resonance Imaging; EEG = Electroencephalography; BOLD = Blood Oxygen Level Dependent; LOO – leave-one out; LOOCV – Leave One Out Cross Validation.

We hypothesized that the reconstructed VS-EFP signal would predict BOLD activity of the VS, even when applied to an independent dataset not used for deriving the model. Furthermore, as the VS forms part of a reward circuit (Haber and Knutson, 2009), we expected that the VS-EFP signal modulation would correlate with additional interrelated regions of the reward network, such as the ventral medial prefrontal cortex (vMPFC). In terms of the EFP model specificity, we expected that the association between the VS-EFP signal and the BOLD activity in the VS would be higher than when compared to an EFP of a different non-reward-related control region. Lastly, in terms of functional validity, we expected that the VS-EFP signal would be modulated by musical pleasure. We further conjectured that the VS-EFP signal modulation would be associated with VS-BOLD signal modulation under different reward contexts, indicating general reward process representation.

## 2. Materials and methods

The project included two experiments with a total of 31 healthy participants; the first aimed to develop the model (termed here *modeling study cohort*) and the second aimed to externally validate the model (termed here *validation study cohort*). The two groups were comprised of different individuals. The studies were conducted at the Tel Aviv Sourasky Medical Center and Tel Aviv University and were approved by the Tel Aviv Sourasky Medical Center and Tel Aviv University ethics review boards. All participants gave their written informed consent. In both studies, the participants underwent the same experimental procedure, and their data were acquired and processed using the same pipeline.

### 2.1 Participants

The modeling cohort included 17 participants (9 females, mean age 25.4 ± 4.56y); and the validation cohort, 14 participants (8 females, mean age 23.9 ± 2.1y). The sample size of the modeling cohort was determined based on previous studies (Keynan et al., 2016) showing SD=0.54 for the statistic of interest; 16 datasets are needed to guarantee a 0.8 power of detecting a (positive) 0.3 average correlation between EFP and fMRI-signal within the target ROI, with a 0.05 chance of falsely finding a non-null correlation. All participants met study participation criteria: (1) they were healthy, right handed, and had normal hearing and normal or corrected-to-normal vision; (2) they met the MRI safety criteria and, (3) they had normal hedonic responses to music, as indicated by a Hebrew-translated version of the Barcelona Music Reward Questionnaire (BMRQ; Mas-Herrero et al., 2013) with a score of 65 or more. Participants were recruited through advertisement and underwent screening via email to assess their eligibility, in terms of MRI safety and study participation criteria.

### 2.2 General Procedure

Participants came to the laboratory twice. The first meeting included a one-hour long behavioral session, which was conducted for the purpose of stimulus selection and included providing their subjective rating of different musical excerpts (see details in the fMRI tasks section 2.2.1 below). During the second session, which was approximately three hours long and took place between 7 and 84 days following the first session (mean = 41 ± 20 days), participants underwent simultaneous EEG/fMRI recordings, immediately followed by a behavioral session outside the scanner. During scanning, participants were presented with a pleasurable music listening task in two separate runs. The datasets from this task served for the EFP-model extraction in study 1 (the modeling cohort) and for its external validation in study 2 (the validation cohort). Twenty-eight participants from both cohorts also completed a monetary reward related task (the MID task) during the EEG/fMRI scanning, in-between the two music runs. Data from this task was used for functional validation analysis. The scanning was immediately followed by a behavioral session outside the scanner that included continuous pleasure ratings of the musical materials. EEG/fMRI data from nine music listening datasets (hereby termed runs) in the modeling cohort and from five runs in the validation cohort were not included due to noisy EEG signal or excessive head movements (>2 mm). One music listening dataset from the validation cohort was further removed as the participant reported to have fallen asleep. Hence, valid EEG/fMRI data were available for 25 runs gathered from 14 different participants in the modeling cohort, and for 23 runs gathered from 12 participants in the validation cohort (modeling: *M*_age_ = 25. 86 ± 4.7; 8 females; validation: *M*_age_ = 23.75 ± 2.16; 8 females). EEG/fMRI data from two additional participants were not included in the analyses of the music-task reward-related modulation due to insufficient (*N* = 1, modeling cohort) or missing continuous rating data (*N* = 1, validation cohort). Data from ten participants in the MID task were excluded due to noisy EEG signal or excessive head movements, resulting in a dataset from 18 participants from both cohorts (*M*_age_ = 24.87 ± 4.11; 12 females).

#### 2.2.1 fMRI Tasks

##### 2.2.1.1 Pleasurable Music Listening Task

This task, which is depicted schematically in Fig. 2a, included passive listening with eyes closed to pleasurable or neutral musical excerpts, and was followed by a separate behavioral test outside the scanner in which the participants continuously rated each musical excerpt for their pleasure level on a scale of 1 (no pleasure) to 4 (peak pleasure). A total of eight excerpts, lasting three minutes each, were presented in two runs, lasting 15 minutes each. Each of the runs consisted of four musical excerpts that were individually tailored: half of the excerpts were self-selected pleasurable excerpts, whereas the other half were rated as neutral and obtained from other participants’ selected excerpts. This counterbalancing ensures that across the entire experiment, there were no consistent acoustical differences in the items considered pleasurable or neutral (Blood and Zatorre, 2001; Salimpoor et al., 2011). It further ensures that the musical materials are completely different across the modeling and validation study cohorts. The individually tailored musical pieces were selected based on a previously established procedure (Blood and Zatorre, 2001; Salimpoor et al., 2011). This approach ensures maximal hedonic responses from each person. Briefly, the musical materials were selected based on an experimental session (during their first visit to the lab) that took place prior to scanning, during which they listened to music and provided their subjective ratings of pleasure (see supplementary materials for details). The musical materials were diverse of different genres, the most common of which were alternative/indie, pop, rock, Israeli music and dance/electronic. The 3-minute music presentation was interleaved with 45 seconds of silence and was delivered using Presentation® software (Neurobehavioral Systems, Inc., Berkeley, CA, www.neurobs.com). During scanning, participants listened to the music through MR-compatible headphones (50-15,000 Hz frequency response) with about 25 dB of passive gradient noise attenuation (Optoacoustics, Israel; for further details about the design, see supp. Materials).

**Fig. 2.**
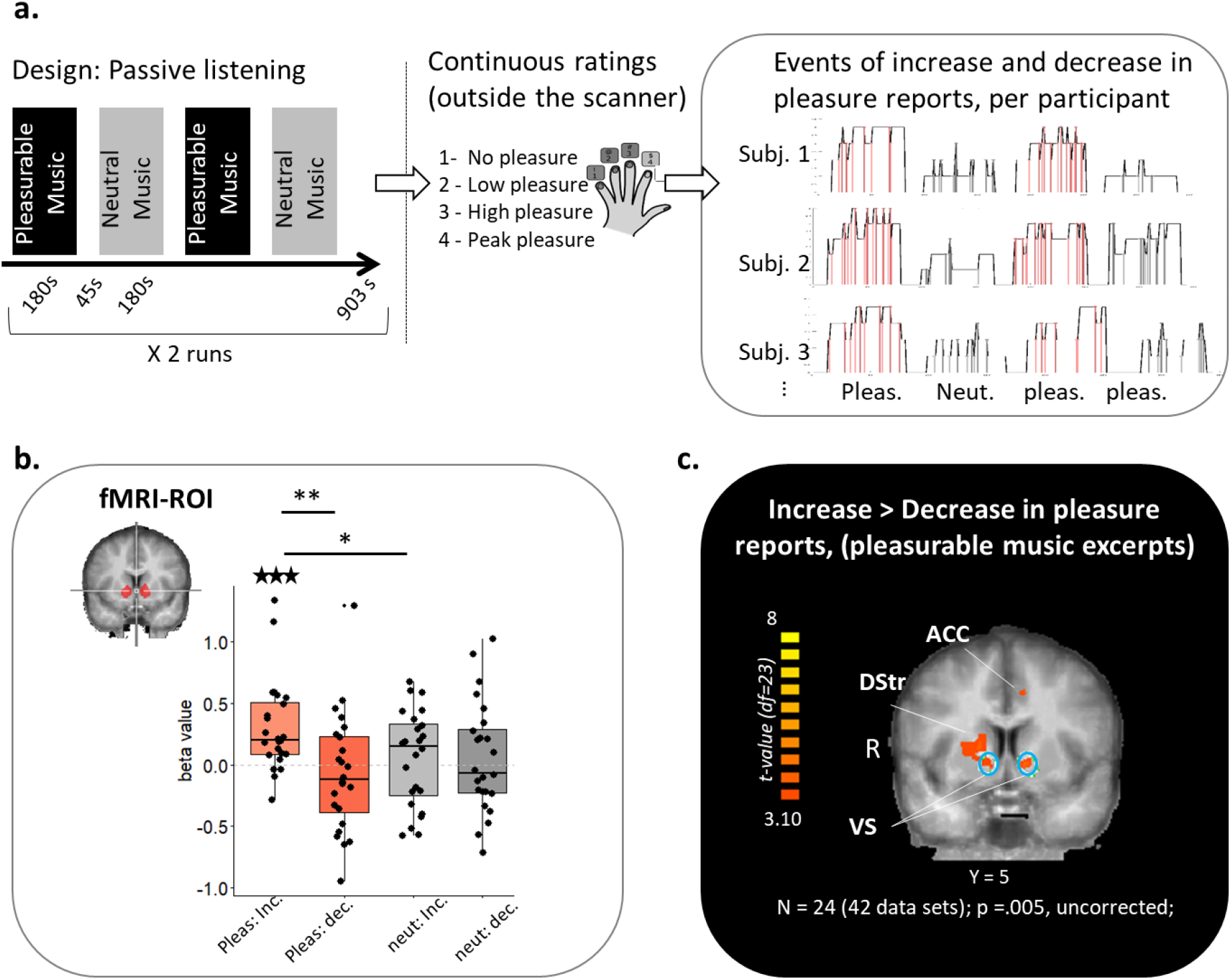
VS BOLD activation modulation by musical pleasure. To induce a reward-related context, EEG-fMRI data was amassed while participants underwent a pleasurable music listening task. **a**. <u>Task design</u>: The task included passive listening to 8 musical clips (four neutral and four pleasurable) inside the scanner, followed by a continuous rating session of the same musical pieces outside the scanner. Using continuous rating, individual increases and decreases in reported pleasure were identified per condition. **b**. <u>ROI analysis</u> of the extracted BOLD data from the bilateral VS across both studies revealed functional selectivity to moments of an increase over a decrease in pleasure ratings while listening to pleasurable music. The boxplots represent the regression coefficients per condition. Paired comparisons: * p<.05; ** p<.01; *** p <.005; Comparison to zero: ★ ★ ★ p <.001. c. <u>Whole brain analysis</u> depicting the transient pleasure increase vs. decrease effects is provided to highlight the associated regions. The target ROI is circled in cyan. Abbreviations: subj. = subject; VS = ventral striatum; ACC = Anterior Cingulate Cortex, DS = Dorsal Striatum; Pleas. = pleasure; neut. = neutral; inc. = increase; dec. = decrease;

##### 2.2.1.2 Monetary Incentive Delay (MID) task

We used a modified version that has been developed and used by Baker and colleagues (Baker et al., 2017). At the beginning of each trial, participants were presented with an anticipation cue that informed them of whether they were expected to gain money in this round or not (monetary gain vs. control conditions). Following the presentation of this cue (500 ms) and a variable anticipatory delay period (jittered around 2,500 ms), the participants were presented with a simple geometric shape and were requested to press a button once they believed that a second had elapsed since its presentation. Following participants’ response and an additional short delay, the feedback – whether they hit or missed the time-estimation goal – was reported on the screen. In the gain condition, a feedback cue notified them whether they won 0.5 ILS (approximately $0.15) or no money (0 ILS) during that trial depending on whether they hit or missed. In the control condition, a visual cue notified whether they hit or missed (using a ‘v’ or ‘x’ mark, respectively). Each of the conditions (gain or control) was presented approximately 60 times in a pseudo-randomized manner. Time estimation thresholds for hits and misses of each trial type were set using an adaptive algorithm to allow the subject to win approximately 60% of the time. As a result, participants were paid an extra sum of up to 10 ILS (approximately $3). Participants were given instructions and underwent a practice run before entering the scanner to familiarize them with the task and to collect baseline data to set the threshold for the initial hit/miss ratio.

#### 2.2.2 MRI data acquisition

Structural and functional MRI scans were performed in a 3T Siemens MAGNETOM Prisma scanner (Siemens, Erlangen, Germany) with a 20-channel head coil. *Anatomical scans* consisted of 3D T1-weighted MPRAGE sequences with a 1 mm iso-voxel to provide high-resolution structural images. *Functional scans* were performed in an interleaved top-to-bottom order, using a T2*-weighted gradient echo planar imaging pulse sequence (TR/TE = 2620/30 ms, flip angle = 90°, 64 × 64 matrix, FOV = 192 × 192 mm, 43 slices per volume with 3 mm thickness and no gap). Positioning of the image planes was performed on scout images acquired in the sagittal plane. A total of 345 volumes were acquired for each of the music listening runs, and between 278 and 332 volumes were acquired for the MID runs. Functional scans of two participants for the MID session were slightly shorter (166 to 237 volumes), due to technical limitations.

#### 2.2.3 Simultaneous EEG recording

EEG data were acquired using a battery-operated MR-compatible BrainAmp-MR EEG amplifier (Brain Products, Munich, Germany) and the BrainCap electrode cap with sintered Ag/AgCl ring electrodes, providing 30 EEG channels and 1 electrocardiogram (ECG) channel (Falk Minow Services, Herrsching-Breitbrunn, Germany). The electrodes were positioned according to the 10/20 system, with a frontocentral reference. The signal was amplified and sampled at 5 kHz and was further recorded using the Brain Vision Recorder software (Brain Products, GmbH, Gilching, Germany).

### 2.3 Data analysis

#### 2.3.1 Preprocessing

##### 2.3.1.1 The fMRI preprocessing

which was done using Brain-voyager QX (Brain Innovation, Maastricht, The Netherlands), included slice timing correction, motion correction using sinc-interpolation, and high-pass filtering of three cycles per scan. Each functional dataset was then manually co-registered to the corresponding anatomical map and incorporated into a 3D dataset via trilinear interpolation. The obtained functional data were then normalized into Talairach space and were spatially smoothed using a Gaussian kernel (isotropic 4-mm FWHM).

###### 2.3.1.1.1 The EEG preprocessing

was conducted using the BrainVision Analyzer software (Brain Products, GmbH, Gilching, Germany). Preprocessing included (1) MR-gradient artifact removal; (2) down-sampling to 250 Hz; (3) low-pass filtering to 70Hz; (4) cardio-ballistic artifact removal using semi-automatic R peak detection, followed by a correction based on the subtraction of an averaged artifact template (Allen et al., 1998); (5) high-pass filtering to 0.75Hz; and (6) notch filtering of 33Hz (31-35Hz) was further applied to account for a periodic scanner-related noise of that frequency.

#### 2.3.2 fMRI statistical analyses

The analysis of the music and the MID tasks were performed according to the random effects general linear model, as implemented in BrainVoyager QX software (Brain Innovation, Maastricht, The Netherlands). For the *pleasurable music task*, eight task-related regressors were included in the model. The pleasurable and neutral music conditions were modeled time-locked to five seconds after the onset of each excerpt. The onsets (first 5 seconds) were also specified in the design matrix as separate regressors. The reward-related responses to music were modeled based on the continuous ratings, which were provided following scanning and were synchronized offline with the scan timing. To account for a response delay of the rating with regards to the musical events (Sloboda and Lehmann, 2001), the rating onsets were shifted by 1 TR (earlier). Responses were divided into moments of increase or decrease in reported pleasure, as events time-locked to the moments at which participants pressed the button, to indicate a positive or negative change in their rating, respectively, per musical condition (i.e., pleasurable or neutral; see Fig. 2a for illustration). *For the MID task*, eight task-related regressors were included in the model. Each of the experimental conditions (anticipation, response, feedback—hit, feedback—miss) per type of trial (monetary gain or control) were modeled, time-locked to the moment at which the cue appeared on the screen. In both tasks, the task regressors were subsequently convolved with the canonical two-gamma hemodynamic response function (using the spm_hrf.m function). Furthermore, six head-motion realignment parameters and mean signal in the white matter were added to the design matrix to account for motion and other non-neural related variance. This model was then submitted into a second-level random effects analysis, to assess the group effects of specific contrasts (e.g., difference between the increase and decrease in pleasure response in the pleasurable music trials, or the difference between hit and miss during the monetary gain or control trials in the MID task). To avoid assumptions about the distribution, statistical tests for the second-level region of interest analysis were performed using non-parametric tests (wilcox.test function in R).

#### 2.3.3 Model extraction and validation

A schematic description of the model extraction and validation process is provided in Fig. 1. The procedure includes: 1. *extraction* of the BOLD response variable from the VS target, and construction of the EEG feature space; 2. *computing of the prediction model* using machine learning to find the combination of electrodes, band-limited power, and time delays that predict the concurrently acquired VS-BOLD signal; and 3. *model evaluation* using a series of analyses that assessed the model’s: a. *sensitivity*, by correlating between the VS-EFP and the BOLD of the VS (ROI analysis) and of additional regions (whole-brain analysis); b. *neuroanatomical specificity*, by comparing the VS-EFP model to a prediction model of another functionally distinct region in the premotor cortex; and c. *functional validity*, by examining if the VS-EFP is modulated by musical reward and whether its prediction of VS-BOLD activation generalizes to another reward-related context.

##### 2.3.3.1 Step 1: Constructing the targeted VS-fMRI signal and EEG feature space for its modeling. *fMRI*

To prepare the predicted BOLD signal, an ROI in the bilateral VS was demarcated. To ensure functional relevance to reward processing, and to avoid circularity, the VS ROI was defined independently from the present data, based on a mask generated from a Neurosynth (http://www.neurosynth.org/ (Yarkoni et al., 2011) map depicting the meta-analysis of the term reward (association test map, threshold = 14.5), which was converted into a binary mask (0/1). The ROI encompassed R+L VS (upper-left inlet in Fig. 1). For every run and participant, one time-series of VS-BOLD activation was constructed using the average activation of all the voxels within this ROI mask. To account for movement-related noise within the extracted BOLD signal, six head-motion realignment parameters were regressed out of the resulting time course, using linear regression. The resulting BOLD signal was normalized to z-scores (zero mean and one standard deviation) and up-sampled to 2 Hz, which corresponds to the final sampling rate of the EEG features (see subsection 2.3.3.1. EEG).

###### EEG

To prepare the EEG feature space, we first accounted for possible artifacts due to head movements, eye blinks, etc., by applying an additional preprocessing step for the automatic detection and removal of non-stationary components in each band, using the analytic approach for Stationary Subspace Analysis (SSA; Hara et al., 2012). This approach includes the analytic separation of the data into stationary and non-stationary sources (i.e., components per frequency band). Using this approach, a component is considered an ‘outlier’ and analytically removed if its associated eigenvalue is larger than a threshold, *P*_50_ + 5 · (*P*_75_ − *P*_25_), where *P*_*i*_ stands for the i^th^ percentile. The resulting time-series were further submitted to an outlier removal procedure, whereby values exceeding the Median Absolute Deviation of five were replaced with a value that was linearly interpolated from adjacent data points in the time series (Leys et al., 2013). We next represented the preprocessed EEG time series in the time-frequency domain in each channel, extracting the log-power of eight frequency bands from the time series of each channel using the function *bandpower*.*m* running on Matlab (2017a, Natick, Massachusetts: The MathWorks Inc.). The band power estimation was performed in sliding windows of 1 [sec] and an overlap of 0.5 [sec], resulting in a time-series with the sampling rate of 2 Hz. The division into bands followed the known division into the EEG frequency bands, as follows: [0-2; 2-4; 4-8; 8-12; 12-16; 16-20; 20-25; 25-40]. Then, to account for the hemodynamic response in the fMRI data, we added time delayed versions of each feature in steps of 0.5 seconds up to 30 seconds, hence generating 60 shifted time-series per band and channel. Finally, the resulting time-series were normalized into z-scores, leaving each frequency with a mean of zero. This feature extraction step resulted in a multidimensional normalized EEG feature space for each time point (T) that included Frequency bands (FQ) X Delays (D) X Electrodes (E) and was used to predict the BOLD activity in the VS (at time point T) during step 2. The data entered into the model-generation procedure contained the entire set of time points of all runs (i.e., 25 runs from 14 participants).

##### 2.3.3.2 Step 2: Modeling of the VS BOLD signal using the EEG features space

The EEG modeling approach seeks to identify a group of electrodes, frequencies, and time-delays in the EEG that, using a weighted linear combination model, predicts the simultaneously acquired VS-BOLD signal at each time point T. Given the high dimensionality of the EEG data, and the highly correlated nature of its features, we applied a partial least square (PLS) regression (de Jong, 1993). The model was trained in two main steps. The first step involved a grid search for selecting (the identity of) a group of up to ten electrodes to be used in the model; and the second step, given the selected electrodes, involved a grid search for selecting the optimal number of PLS components (with a maximum of 10) to predict BOLD data from the EEG channel’s group feature space. Extending the previous EEG modeling approach (Meir-Hasson et al., 2016, 2014), here we select a group of channels, rather than a single channel, to construct the prediction model. The channel selection is performed using a grid search, from one to ten channels. The identity of each group of channels was selected by adapting the group LASSO approach (Lassoed principal components) of Witten and Tishibirani (2009). This approach involves the fitting of a PLS model with a penalty on groups of coefficients (each group corresponds to a channel), thus searching for a parameter that optimizes a selected group of channels and penalizes the rest (L21; PLS-1 with LASSO). The optimization in this step is based on the first PLS component. More specifically, in this procedure we aimed to solve the following optimization problem:

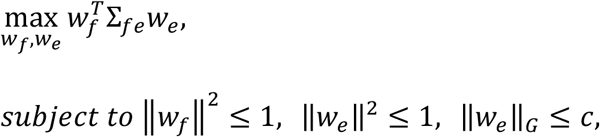

where Σ_*fe*_ is the covariance matrix of the concatenated fMRI and EEG features time series; *w*_*e/f*_ are the weights of EEG and fMRI, respectively, whose aim is to maximize the covariance between the EEG and fMRI component; ‖·‖_*G*_ is the group lasso penalty; and c is a parameter that controls the group lasso penalty. Since the fMRI data contains a single time series, we set *w*_*f*_ = 1 and optimized *w*_*e*_. In our implementation, we searched for the value of *c*, such that the desired number of channels is selected.

With the selected group of channels (from the output of the previous stage, groups ranging from 1 to 10 channels), we turned to a second grid search to select the optimized PLS component number out of a maximum of ten. This step was applied by fitting the PLS model (Matlab function *plsregress*.*m*, applying SIMPLS algorithm, (*78*)), each time with the group of selected channels. This separation into two steps was undertaken to avoid the group lasso penalty bias, as well as for computational efficiency.

To estimate the best values for these parameters, we used a grid search and the cross-validation method. For cross-validation, we used the leave-one-run-out cross-validation (LOOCV) method, wherein in each fold, one of the runs is left out for test, and the model is trained on the rest. These two steps resulted in a performance matrix of number of channels by number of PLS components (i.e., 10 * 10) per each of the N folds (i.e., number of runs across participants) of the external LOOCV. The performance was indexed as the average correlation between the model’s output and the fMRI BOLD signal in the VS region of interest across the leave-one-out datasets. The selected model, termed the VS-EFP model, was the one with maximal average performance across the channel and component numbers.

##### 2.3.3.3 Step 3: Model evaluation and validation and statistical analyses

The modeling process resulted in the selection of a model represented as a time-delay × frequency x electrode weight coefficient matrix, (see Fig. 1.5 for a depiction of the model’s coefficient matrix). The weights of this model could now be applied to the EEG feature space (D * FQ * E) at time point T to reconstruct the VS-EFP signal for that time point. The derived VS-EFP signal was submitted to a series of analyses that aimed to assess its sensitivity, neuroanatomical specificity, and functional validity (see details in the model evaluation subsections, 2.3.3.3.(1)-(4)). The model’s performance was initially evaluated using the leave-one-run-out-cross validation approach that was applied as part of the modeling procedure. Importantly, the model’s performance was then assessed using an out-of-sample validation approach applied on the validation cohort. Finally, the functional validity of the VS-EFP signal in probing reward-related processes was examined by testing if it is sensitive to changes in the experience of musical reward and whether its performance generalizes to another reward-related context (the MID task). Specifically, the following validation analyses were performed: (1) **Model sensitivity**. <u>ROI analysis</u>: The model’s performance was indexed per dataset as the correlation between the VS-EFP and the VS-BOLD signals. The VS-BOLD signals were extracted from the same ROI used to derive the EFP model. The likelihood of the obtained correlation was assessed by determining the percentile location of the empirical value within the null distribution of performance values. The null distribution was reconstructed using a phase-randomization permutation approach (Lerner et al., 2011), by repeating 1,000 times the same calculation for estimating the performance, with the important exception that the VS-BOLD signal was phase-randomized before the correlation was calculated. <u>Whole-brain analysis</u>: To assess the whole brain sensitivity of the VS-EFP signal, a whole-brain random-effects general linear model analysis of the fMRI data was performed, using the VS-EFP signal time-series as the regressor. The resulting regression coefficients were submitted to a second-level analysis using a one-sample t-test. (2) **Neuroanatomical specificity of the VS-EFP:** random effects (ROI and whole-brain) analyses were conducted to compare between the VS-EFP and an EFP of a functionally distinct region in the premotor cortex, hereon termed premotor-EFP. The premotor-EFP model was derived using the same machine-learning pipeline as described above in steps 1 and 2, with the exception that now the modeling was focused on predicting the BOLD response in the right premotor cortex. This region was defined functionally based on a statistical parametric map that was obtained from the whole-brain contrast of music vs. baseline in the modeling cohort (q(FDR)<.05, *p* < .002; *n* = 16). To avoid multi-collinearity, the two regressors were orthogonalized using the *spm_orth*.*m* function, such that the additional EFP (premotor- or VS-EFP) was orthogonalized relative to the EFP signal of interest (either the VS-EFP or premotor-EFP). (3) **Functional validation of the VS-EFP, reward-related modulation:** To assess whether the VS-EFP signal is modulated by musical pleasure, the VS-EFP time-series were submitted to a two-level random effects general linear model analysis, using the same predictors and procedure that were used for delineating the BOLD response to the reward task (see section 2.3.2, above, for details). To avoid assumptions about the distribution, the second-level analysis was applied using non-parametric statistics. (4) **Functional validation of the VS-EFP, generalization to another reward related context:** Examination of the association between VS-EFP signal and the VS-BOLD activity under a different reward-related context abundantly in reward studies; the MID task (Lutz and Widmer, 2014). This was achieved by assessing the sensitivity of the VS-EFP using a similar procedure as described above in section 2.3.3.3.(1), now applied to the MID task data.

In all the above-mentioned general linear model analyses, the EFP time-series were submitted to an outlier removal procedure that was applied with the *icatb_despike_tc*.m function of the GIFT toolbox (http://trendscenter.org/software/gift/). Six head-motion realignment parameters and mean signal in the white matter were further added to regress out motion and another non-neural-related variance. In the ROI analyses, outlier data points exceeding 3 * interquartile range from the 25^th^ or 75^th^ percentiles were excluded from analysis, and second-level analyses were conducted using non-parametric statistics (i.e., Wilcoxon signed rank test). Finally, correction for multiple comparison was achieved by applying the Benjamini-Hochberg procedure for controlling the false discovery rate (FDR; Benjamini and Hochberg, 1995). The statistical threshold of significance was further set with a minimal cluster-size of five contiguous functional voxels (>125 mm3).

## 3. Results

### 3.1. VS-BOLD activation modulation by musical reward

In the present study, we sought to extract an EEG model of reward-related VS-BOLD activation. To induce naturalistic reward related activation, participants listened to individually tailored pleasurable music (Fig. 2a). Therefore, the first step was to verify that the VS-BOLD signal was modulated, as expected, by musical reward. For this purpose, we conducted a region of interest (ROI) analysis using the same bilateral VS mask that had been used as to generate the VS-EFP. As shown in Fig. 2b, we found an enhanced VS-BOLD response to musical pleasure. Specifically, while listening to pleasurable music, there was higher VS BOLD activation during moments of increase (i.e., a positive change) in pleasure ratings relative to baseline (median β= 0.13; one-sample Wilcoxon signed-rank test; *V* = 258; *p* < .001), which was greater than the response to moments of decrease (i.e., negative change) in rating (paired Wilcoxon signed-rank test; *V* = 228; *p* = .0024), as well as greater than the response to moments of increase in pleasure ratings while listening to the neutral music (paired Wilcoxon signed-rank test; *V* = 196; *p* = .041). Whole-brain depiction of this transient reward-related effect (i.e., increase vs. decrease in pleasure) highlighted additional reward-related regions beyond the VS; namely, the anterior cingulate area (ACC) and dorsal striatum (*p*<.005, uncorrected; Fig. 2c). This analysis demonstrated the functional relevance of the experimental design in engaging the VS region used for modeling.

### 3.2. Characterization of the selected VS-EFP model

The concurrently acquired EEG/fMRI data from the modeling study cohort were used to calculate the VS-EFP prediction model using a machine learning procedure. The VS-EFP prediction model was selected based on a grid search as the model that showed the highest performance; i.e., the highest average correlation between the VS-BOLD and the VS-EFP (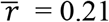; see grid search results in Fig. 1.). The selected model was derived using a group of eight electrodes (C4, F7, F8, T7, T8, P8, TP9 and TP10) and three PLS components. The model weights per time-delay, frequency band and electrode are depicted in Fig. 1.5.

### 3.3. VS-EFP model sensitivity

#### 3.3.1. ROI analysis - Target sensitivity

The first step was to assess the sensitivity of the VS-EFP model in predicting the BOLD signal in the VS ROI that was used for the modeling, relative to a phase-permuted version of the VS signal. This evaluation was achieved by computing the correlation between the reconstructed VS-EFP time-series and the concurrently acquired VS-BOLD and assessing these values relative to a null distribution. The prediction performance of the VS-EFP model was assessed per cohort. Fig. 3a shows the results obtained for the modeling cohort, using the leave-one-out cross-validation approach. Permutation testing based on phase randomization ensured that the mean performance (i.e., correlation) is significantly better than the mean performance of the VS-EFP model in predicting phase-randomized versions of the VS-BOLD (*p* < 0.001). Inspection of the model performance at the individual level further revealed a significantly higher performance than expected by chance (*p* <.05) in 12 out of the 14 participants (and in 20 out of the 25 datasets). Importantly, we next examined if such sensitivity is evident when the VS-EFP model is applied on a new dataset of the validation cohort, which was not used for modeling. Fig. 3c shows the sensitivity testing of the VS-EFP model for the validation cohort (N=12, 23 datasets), revealing an average correlation of 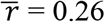, which was significantly better than the phase-randomized null distribution of mean performances (*p* < 0.001) and in a similar range as observed in the modeling cohort. Inspection of the model performance at the individual level with permutation tests further revealed a significantly higher performance than expected by chance in all 12 participants (and in 20 out of the 23 datasets; *p* <.05). Cross-correlation analysis that was computed between the VS-EFP and VS-BOLD further unraveled that VS-EFP prediction is well synchronized with the VS-BOLD in both datasets (maximum correlation is at zero delay, as shown in Fig. 3a,c, right panels).

**Fig. 3.**
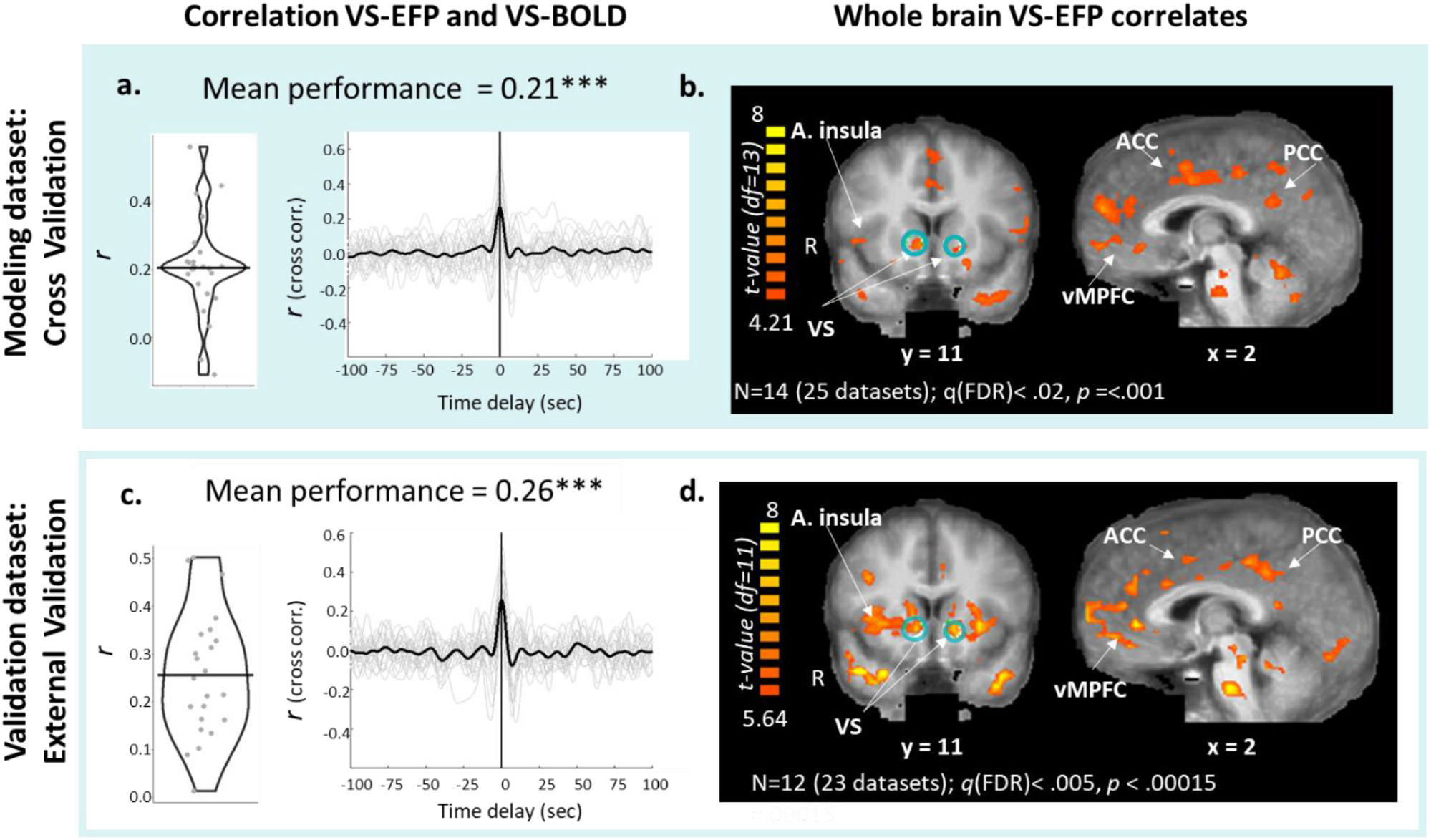
VS-EFP model sensitivity. The sensitivity of the VS-EFP in predicting the BOLD signal within the VS and associated regions was examined by assessing: **a**. <u>Target sensitivity</u>: evaluated as the correlation (left panel) and cross-correlation (right panel) between the time series of VS-EFP and the predicted BOLD signal within the target VS ROI. **b**. <u>whole-brain sensitivity</u>: evaluated by applying a two-level random effects general linear model analysis to highlight the correlates of the VS-EFP at the whole-brain level. These analyses were first carried out on the modeling study cohort using the leave-one-out-cross validation approach. To externally validate the model, the same **c**. ROI and **d**. whole brain analyses were performed on the validation study dataset, an independent dataset of a completely different group that underwent the same EEG-FMRI scanning protocol. The target ROI is circled in Cyan. Abbreviations: vMPFC = ventromedial prefrontal cortex; A. = anterior; PCC = posterior cingulate cortex; STG = superior temporal gyrus; (*** p <.001).

#### 3.3.2. Whole-brain sensitivity – whole-brain correlates of the VS-EFP

Next, we sought to highlight the additional BOLD regional activation that the VS-EFP signal is sensitive to in each cohort. Towards this aim, we applied a whole-brain random effects general linear model analysis of the fMRI using the VS-EFP signal as the predictor, while setting the highest statistical threshold that allows the detection of activity within the VS. This analysis revealed that the VS-EFP signal is correlated with the BOLD activity within the VS at a threshold of q(FDR) < 0.02 (peak Talairach coordinates within the target ROI: R. VS: x = 6, y = 9, z = 1; *t*(13) = 5.91; L. VS: x = -12, y = 11, z = -2; *t*(13) = 4.57; *p*<.001; Fig. 3b), as well as with additional brain regions related to the reward network, including the ventromedial prefrontal cortex (vMPFC), anterior insula, and ACC, and also with additional regions related to more general processes such as the Posterior cingulate cortex (PCC), thalamus, and right posterior superior temporal gyrus.

Application of whole-brain VS-EFP sensitivity testing to the *validation cohort* at a matched statistical threshold with a similar number of voxels (∼80,000 correlated voxels) showed that the VS-EFP signal correlated with the BOLD signal modulation within the VS at a threshold of q(FDR) < .005 (R. VS: x = 9; y = 8, z=1; *t*(11) = 7.71; L. VS: x = - 15, y = 8, z = -2; *t*(11) = 9.03; *p*<.0001; Fig. 3d), along with additional functionally related regions such as the anterior insula ACC, amongst others. Regions that were consistently correlated with VS-EFP signal across the modeling and validation datasets were further highlighted using a probability map, which was constructed from the two statistical parametric maps (Fig. S1; for a full list of the highlighted regions, see Table S1). In both datasets, in addition to the VS, the VS-EFP consistently correlated with functionally related regions such as the vMPFC, anterior insula, amygdala, PCC, and thalamus, as well as with additional regions such as the STG and the cerebellum.

### 3.4. Neuroanatomical specificity of the VS-EFP model

The analyses thus far suggested that the VS-EFP signal is sensitive to the BOLD signal modulation in the VS, as well as in additional functionally related brain regions. This finding raises a question regarding the degree to which the model of the VS is neuroanatomically specific. To address this question, we used an EFP model of the premotor cortex – a functionally distinct region that is also modulated during music listening – as an additional fMRI whole-brain predictor. This inquiry enabled a performance comparison between VS and premotor EFPs in predicting VS-BOLD vs premotor activation. Random effect general linear model analyses were applied within these two target ROIs, using the two EFP-signals as regressors (see materials and methods section 2 for details). This analysis revealed notable target-EFP model specificity in the two cohorts. Specifically, the VS-BOLD modulations were more strongly associated with the VS-EFP than with the premotor-EFP signal modulation in the modeling cohort (paired Wilcoxon test, *V* = 12, *p* = .009, Fig. 4a), and importantly, also in the validation cohort (paired *V* = 3, *p* = .002, Fig. 4b). Similarly, the premotor-BOLD fluctuations correlated more strongly with the premotor-EFP than with the VS-EFP consistently in both datasets (modeling cohort: *V* = 104, p<.001, Fig. 4c.; validation cohort: *V* = 77, p<.001, Fig. 4d; see Fig. S2 for the full comparisons). Whole-brain depiction of these effects was further performed to highlight the spatial distribution of this EFP-specificity in both datasets. The results of this analysis, as depicted in Figs. 4 e-h, revealed a compelling network specificity per EFP model. In particular, it is evident that the VS-EFP relative to premotor-EFP was significantly more correlated with the BOLD activity in the ventral and dorsal striatum and ACC, both in the modeling cohort (Fig. 4e; *p* =.0072, uncorrected) and the validation cohort (Fig. 4f, lower panel; *p* =.001; q(FDR)<.1; for a list of the highlighted regions, see Table S2). Correspondingly, a comparison between the premotor-EFP and the VS-EFP revealed that this motor-centered EFP was significantly more correlated with a different set of regions, belonging to a motor and somatosensory network including the supplementary motor area, the primary motor cortex, and the premotor cortex both in the modeling (Fig. 4g, upper panel; *p* =.0072, q(FDR)<.05) and validation cohorts (Fig. 4h, lower panel; p = .001, q(FDR)< .01).

**Fig. 4.**
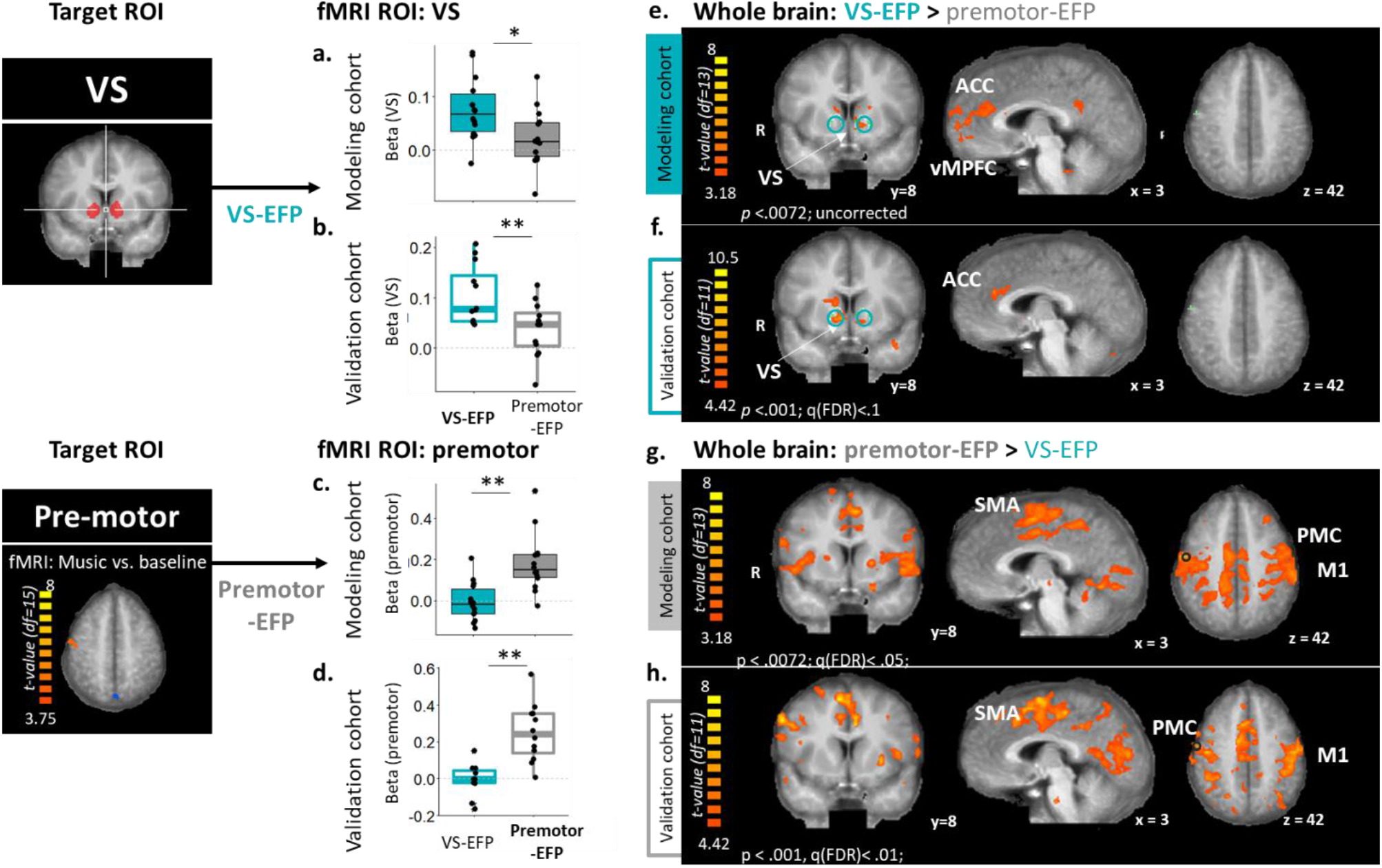
Neuroanatomical specificity of the VS-EFP model; The specificity of the EFP-model (top panel) in predicting the targeted BOLD signal and associated regions was assessed by contrasting the BOLD correlates of the VS-EFP to the BOLD correlates of an EFP of another functionally distinct target region in the premotor cortex, and vice versa (lower panel). The ROI within the premotor cortex that was used for extracting the EFP model was highlighted based on the contrast fMRI map depicting the difference between music vs. baseline conditions in the modeling cohort (n=16; left panel). **(a-d)** <u>fMRI-ROI analysis</u>: a comparison between the prediction of each EFP model and the BOLD activity of the target ROI was carried out separately for the (**a**.,**c**.) modeling and the (**b**., **d**.) validation study cohorts. The boxplots represent the regression coefficients per EFP. (Paired comparisons: * p<.05; ** p<.01). (**e-h**) Whole brain analysis: unique whole-brain correlates of the two distinct EFP models are depicted on coronal, sagittal, and axial views for the contrasts: (e., f.) VS-EFP > premotor EFP and (g., h.) premotor EFP >VS-EFP. The target ROIs are circled in Cyan (VS) or black (premotor). **Abbreviations**: SMA = supplementary motor area. M1 = primary motor cortex. PMC = premotor cortex

### 3.5. Functional validation

#### 3.5.1. VS-EFP reward-related modulation

The next step was to examine if the VS-EFP is also sensitive to a reward context induced by music, in a manner akin to the reward-related engagement demonstrated in the VS-BOLD activation. Towards this aim, we applied random effects general linear model analysis with the VS-EFP as the dependent variable and the various modeled reward-related responses to music that were used in the fMRI analysis as predictors. This analysis revealed a transient VS-EFP response to musical pleasure, as evidenced in the VS-BOLD among the validation cohort (Fig. 5). Specifically, there was an enhanced VS-EFP response during pleasurable music listening in moments of increase in pleasure ratings (Median _β pleasure rating increase_ = 0.24, IQR = 0.32, Wilcoxon one sample signed rank exact test: *V =* 60; *p* =.007). That response was greater than the response to moments of decrease in rating (Median _β decrease pleasure_ = -0.306, IQR = 0.256; paired signed rank comparison: *W =* 59; *p* =.009), as well as greater than the response to moments of increase in ratings during neutral music listening (Median _β increase neutral_ =-0.296, IQR = 0.489; paired signed rank comparison: *W*= 57, *p* = .016). These functional modulations were not evident when examining the leave-one-out EFP models in the modeling cohort. One possible source for this variance is that the observed VS-EFP selectivity to musical pleasure was affected by the extent of reward-related engagement of the VS-BOLD across individuals. To examine this possibility, we tested the correlation between the indices of VS-EFP and VS-BOLD selectivity for increase in pleasure during listening to pleasurable music across both datasets. There was a positive correlation between the selective responses of the VS-EFP and VS-BOLD to pleasurable music during moments of increased vs. decreased pleasure (*r*_*Spearman*_ =.49, *p* < .005), as well as vs. increase in rating during neutral music listening (*r*_*Spearman*_ =.53. *p* < .005).

**Fig. 5.**
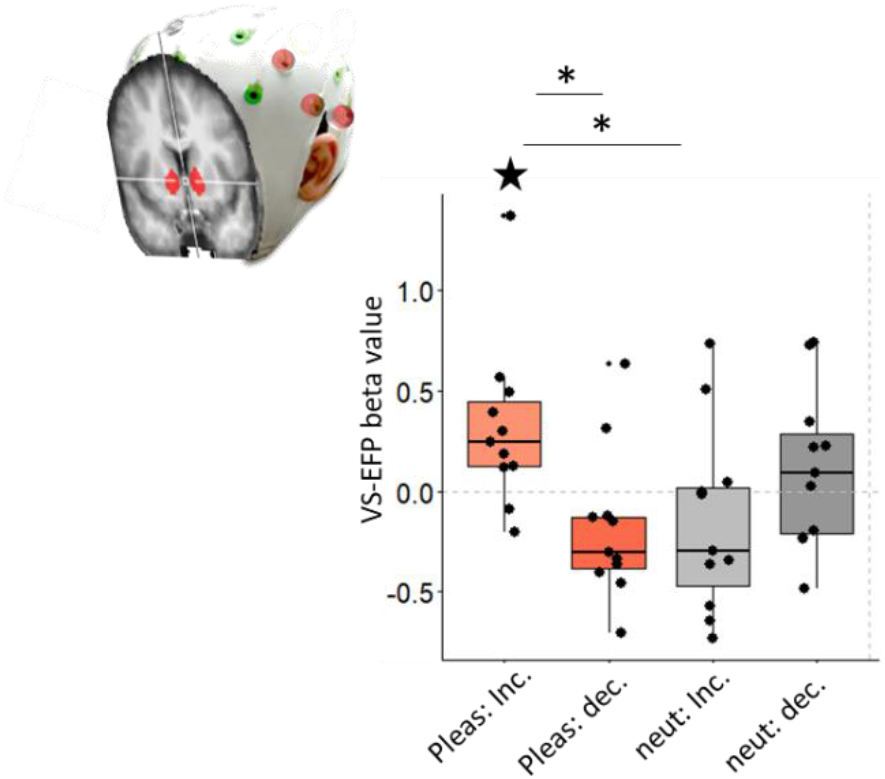
VS-EFP reward-related modulation. To asses if the VS-EFP was functionally modulated by musical reward, a random effects GLM analysis was applied to predict the VS-EFP signal among the validation study cohort dataset (*n*=11). Boxplots represent the resulting regression coefficients per condition, with dots depicting individual values. Paired comparisons: * p<.05; Comparison to zero: **★** p <.05).

#### 3.5.2. Generalization into a different reward context

Finally, we sought to test whether the association between VS-EFP and VS-BOLD activity would be apparent under a different reward-related task and context; namely, the MID task. The average correlation between the VS-EFP signal and the VS-BOLD activity during this task (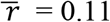 Fig. 6a) was statistically significant, as revealed by using a phase randomization-based permutation testing (*p* < 0.001). Inspection of model performance at the individual level further revealed a significantly higher performance than expected by chance (*p* =.05) in eight of the 18 participants and a performance marginally higher than expected by chance in another four of the 18 participants (up to *p* < .12). A whole-brain random effects general linear model analysis, with the VS-EFP signal as a regressor of interest, further showed that the VS-EFP signal correlated with the BOLD activity of the VS (peak voxel: right: x = 6, y = 12, z = 4, left: x = -9, y = 14, z = 7; N = 18; Fig. 6b). Notably, the VS-EFP signal also correlated with activity in additional reward-relevant brain regions, including the VTA, anterior insula, ACC, thalamus, and amygdala, as well as in additional regions such as the visual cortex (for a full list of the highlighted regions, see Table S3).

**Fig. 6.**
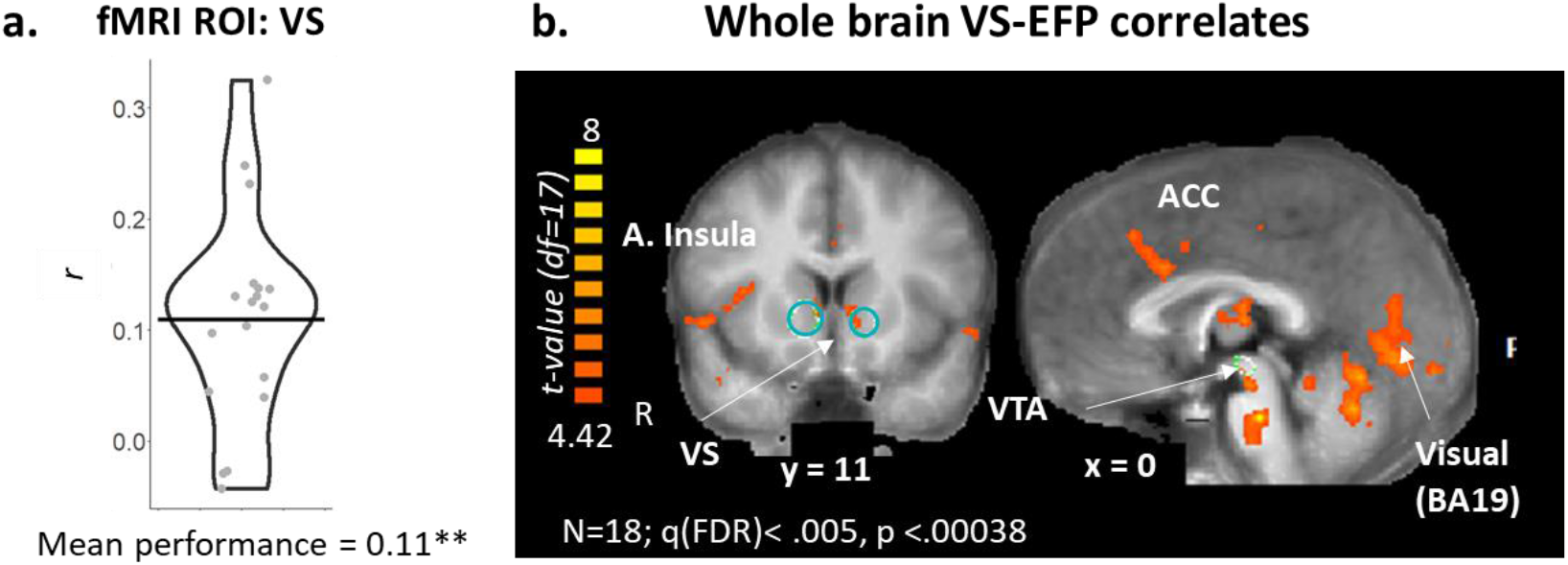
Functional validation; generalization into a different reward context. Examination of VS-EFP performance (sensitivity) in dataset amassed under a different reward-related context induced by the MID task. **a**. <u>VS-EFP model performance</u>: assessed as the average correlation between the time series of VS-EFP and the predicted BOLD signal in the target VS. **b**. <u>Whole-brain correlates of the VS-EFP</u> within this different context revealed a significant correlation between the VS-EFP and BOLD activation in the VS, as well as additional functionally related regions such as ACC, anterior insula, the Ventral tegmental area (VTA), and visual areas.

## 4. Discussion

The aim of this study was to develop and validate an accessible and affordable probe of neural activation related to reward processing in the VS – a deeply located core node of the brain’s reward circuitry. Using an fMRI-informed EEG modeling approach, we identified a particular spatial-temporal-spectral EEG representation that is predictive of the concurrently acquired fMRI activity in the VS while responding to rewarding stimuli (i.e., VS-EFP). A series of validation analyses revealed the VS-EFP model to be correlated significantly with the VS-BOLD signal across individuals, both in the modeling cohort and, importantly, in an out -of-sample validation cohort. We were also able to show the network specificity of the VS-EFP signal by contrasting it to an EEG model of another region derived with the same analytic pipeline. To close the loop between VS-EFP and reward processing, we demonstrated the functional relevance of the VS-EFP signal through its correlation with reported musical pleasure within the validation cohort.

Further alluding to the functional robustness of the VS-EFP model, we found that it also correlated with the VS-BOLD signal under a different context of monetary rewards, indicating that it captures some core aspects of reward processing beyond a particular reward type that used for its development. Jointly, these findings provide compelling evidence of the relevance of the VS-EFP model to serve as a scalable (accessible and affordable) probe of reward-related brain activation.

### 4.1. Model development and validation

In line with our primary assumption, we showed that the VS-EFP signal predicted the BOLD-VS activity, even when tested in an out-of-sample validation dataset. Although it was developed using a leave-one-out cross-validation approach, which could lead to over-fitting, the one-class model performed well in datasets that were not used for the modeling, thus indicating its stability.

The idea of developing a common, generic EEG model instead of an individually tailored EEG model of localized brain regions was introduced before (Meir-Hasson et al., 2016). In this initial work, which focused on the amygdala, they used hierarchical clustering to highlight a subgroup of participants from which they derived a single model. They demonstrated that this common model provides better accuracy than the individual model when applied to a new dataset or participant. Furthermore, the amygdala-EFP model was shown to predict amygdala-BOLD activation in a separate validation cohort (Keynan et al., 2016). Here, we followed the rationale of this approach, but instead of preselecting a subgroup of datasets, we applied regression across the entire sample a-priori. The motivation was to enhance the generalization capabilities further.

The VS-EFP was developed and validated within the context of musical pleasure. We probed this experience using individually tailored musical materials, which differed across participants and cohorts, thus further ensuring that the VS-EFP model does not depend on any particular stimulus, sound or acoustic features and is generalizable across a wide variety of musical materials. Supporting this generalization further, the correlation of VS-EFP and VS-BOLD was also evident in response to monetary rewards during the MID task (Fig. 6). Jointly, these observations provide compelling support for the utility of the one-class VS-EFP model in probing reward processes in different subjects and under different reward contexts.

### 4.2. VS-EFP model network correlates

A whole-brain inspection revealed that the VS-EFP signal correlates with several reward-related regions beyond the VS, such as the ACC and the insula, suggesting that this signal captures a VS-related network. Interestingly, such network manifestation was evident within the context of music listening (Fig. 3) and monetary reward (Fig. 6). In addition to these consistently highlighted regions, context-dependent VS-EFP signal correlates were evident elsewhere in the brain. Specifically, during music listening, VS-EFP signal correlated with a network of functionally related regions, such as the PCC and other areas probably associated with the task’s modality; for example, the right posterior STG (BA 22). During the MID task, the VS-EFP signal correlated with a network of functionally related regions, such as the anterior insula, VTA, and additional areas such as the visual cortex and the somatomotor regions, probably associated with the task’s modality. Hence, while a core set of reward-related regions was related to the VS-EFP signal in both tasks, additional correlated areas were task-specific.

This observation resonates with the findings of recent meta-analyses that compared responses to distinct types of reward, such as food and music, which represent primary and secondary rewards with varying levels of abstraction (Mas-Herrero et al., 2021; Sescousse et al., 2013). In these studies, while both reward types engaged mesolimbic regions – including the VS, vMPFC, and the insula – additional activations were modality-specific, such as the STG for musical reward and the amygdala for food reward (Mas-Herrero et al., 2021), or anterior insula for erotic and food more than for monetary rewards (Sescousse et al., 2013). The highlighted common and distinct reward-correlates in these meta-analyses were interpreted as pointing to a core reward network that is commonly recruited during hedonic reactions, though via specific input pathways depending on the reward type or sensory modality. Our observation of distinct VS-EFP correlates under different reward contexts may be interpreted along this line, as reflecting task- and modality-specific pathways through which reward is accessed or processed. Our subsequent finding following a direct comparison between the VS-EFP signal and another EFP, of the premotor cortex, further supports this suggestion. Specifically, a direct contrast between the two anatomically distinct EFP models revealed marked network specificity; while the unique functional correlates of the VS-EFP signal were more restricted to a well-defined mesolimbic network, centered on the striatum and ACC, the correlates of the premotor EFP were in regions of the motor network.

### 4.3. Possible applications of the VS-EFP model

What possible applications could be developed for accessible probing of VS-related activity? Tracking reward-related mesolimbic activity with EEG, by considering its spectral, spatial, and temporal features, may pave the way for investigating various facets of reward processing under more ecological settings, and in environments that are more diverse. For example, investigation of group dynamics may be enabled using EEG hyper-scanning in social contexts, such as during music performances (e.g., Acquadro et al., 2016).VS-EFP signal recordings may also facilitate studies on cross-cultural differences in mesolimbic processing in remote communities with no neuroimaging access (Egermann et al., 2015). Portable or wearable EEG may further extend the investigation to real-life tasks requiring mobility (e.g., sports, playing an instrument, riding a bicycle, etc.;(Scanlon et al., 2019)), or to home-based monitoring(Shustak et al., 2019). Indeed, our observations that the VS-EFP probe was modulated with transient changes in behavioral reports of musical reward in the validation cohort, although requiring further replication, allude to the potential of using the VS-EFP as a scalable probe for tracking reward-related VS function with EEG in various contexts. A parallel demonstration using the EFP modeling approach for accessible probing comes from previous work that examined responses to an anger-provoking film by showing that an EFP of the amygdala tracked ongoing changes in subjective anger experiences (Lin et al., 2017). Notably, such state-induced amygdala-EFP modulations further predicted the levels of post-traumatic stress symptoms a year after stress exposure during military service experiences (Lin et al., 2017).

Beyond its possible use in basic scientific research, the clinical potential of such a probe is noteworthy. Reward circuit disturbances, with the VS at its heart, have been associated with debilitating neuropsychiatric symptoms such as anhedonia (Pizzagalli, 2014), apathy (Levy and Dubois, 2006), and addiction (Luijten et al., 2017) in a range of disorders, such as Major Depression Disorder (Rizvi et al., 2016),Parkinson’s disease (Kirschner et al., 2020), and Substance Use Disorder (Luijten et al., 2017). Hence, monitoring of this circuitry in real-life settings, as well as neuromodulation of this circuitry, may be highly valuable for treating such disturbances. In recent years, there has been a growing interest in using neurofeedback – a non-invasive technique for self-neuromodulation based on a closed-loop for non-pharmacological brain-guided treatment (Lubianiker et al., 2019; Thibault et al., 2018; Trambaiolli et al., 2021). Neurofeedback enables individuals to modulate a specific brain area or a circuit using auditory or visual feedback based on the measurement of their-own brain activity or connectivity (Lubianiker et al., 2019). A variety of brain-recording techniques, such as EEG (Gruzelier, 2014), fMRI (Sulzer et al., 2013a) and even intracranial EEG (Yamin et al., 2017), can be used to guide neurofeedback training procedures. Evidence suggests that individuals indeed can volitionally regulate their regional neural activation, including in deep brain regions such as the VS (Greer et al., 2014; Kirsch et al., 2016; Li et al., 2018) or VTA (Kirschner et al., 2018; MacInnes et al., 2016; Sulzer et al., 2013b) via real-time fMRI (Thibault et al., 2018). Moreover, using real-time fMRI, learnt NF-modulation of deep-brain regions, as the amygdala, was effective in reducing depressive symptoms in Major Depressive Disorder (Linden et al., 2012; Mehler et al., 2018; Trambaiolli et al., 2021; Young et al., 2014). This finding suggests that similar modulation of specific reward-related processes may be beneficial (Lubianiker et al., 2019). Yet despite its promise, to date, only one study examined the potential of mesolimbic self-modulation with real-time fMRI for clinical purposes (Kirschner et al., 2018). One possibility for the scarcity of neurofeedback solutions related to reward circuitry is that the utility of real-time-fMRI-neurofeedback for repeated training is limited due to immobility, high cost, and extensive physical requirements. Implementing the VS-EFP signal together with a neurofeedback procedure may allow for accessible yet process-specific training. A similar approach has been taken using the amygdala EFP to examine the effects of repeated training sessions targeting the self-regulation of this signal (Meir-Hasson et al., 2016). Results from a series of validation experiments of this method with both healthy participants (Keynan et al., 2019, 2016) and clinical populations (Post Traumatic Stress Disorder; *62*, Fibromyalgia; *63*) indicated that individuals are able to learn how to down-regulate the amygdala-EFP, and that such training is associated with marked neurobehavioral changes related to amygdala down-regulation, with significant clinical outcomes. Applying a similar process-based neurofeedback approach (Lubianiker et al., 2019) with the VS-EFP signal while interfacing with pleasurable music – a strong driver of reward-related networks – may prove to be a promising avenue for training individuals to self-regulate this cardinal circuit precisely.

### 4.4. Limitations

We acknowledge that despite accumulating evidence pointing to the feasibility of detecting subcortical electrophysiological activity, including striatal, with scalp EEG (Fahimi Hnazaee et al., 2020; Foti et al., 2011; Seeber et al., 2019), the possibility of accessing activity in deep subcortical regions such as the VS from the scalp is still debated (Cohen et al., 2011; Puce and Hämäläinen, 2017). Hence, a more conservative assumption is that the VS-EFP signal is also driven by cortical sources that correlate with this subcortical area, reflecting a core network related to the VS rather than the activation in the VS *per se*. This suggestion corresponds with the observation that the VS-EFP signal correlates included additional relevant brain regions (Figs. 3-4). Future studies could test this assumption using simultaneous recordings of depth electrodes from the VS and scalp EEG (Seeber et al., 2019) or magnetoencephalography, which can detect deep sources with appropriate modeling (Freeman et al., 2009).

Additional sources contributing to the VS-EFP signal prediction may be non-neuronal and result from physiological or movement noise. One possible solution for reducing such contribution, which was applied here, is to regress out possible nuisance factors. An additional solution that can be tested in future work is to regress out EFPs of another region of no interest.

The marked network specificity that was highlighted in this study by contrasting the VS-EFP signal and premotor-EFP reveals the potential of the latter suggestion.

A further caveat is that the modeling performance, assessed as the correlation between VS-BOLD and VS-EFP signal, was of moderate size on average across participants (mean *r* = 0.25 in the validation cohort). The magnitude of such correlation is comparable to the correlation magnitude reported for the previously developed and validated amygdala EFP model (Keynan et al., 2019), as well as in additional multi-modal studies that examined the correlation between two modalities (Seeber et al., 2019; Wirsich et al., 2021). This moderate relation between the signals might raise the possibility that the EEG and fMRI may probe partially non-overlapping neuronal populations (Freeman et al., 2009), or that the VS-BOLD may partially reflect non-neuronal but meaningful contributions, such as vasculature (Bright et al., 2020).

Finally, the validity of the VS-EFP signal was demonstrated in an out-of-sample validation, despite a limited number of participants, which suggests the result is robust. However, further replication of this association in other cohorts of individuals with different characteristics and in more varied experimental contexts will still be necessary, especially for the functional validation. Future development may also apply a larger dataset for developing the model and the use of advanced algorithms for modeling, such as deep learning. Another improvement of the modeling may incorporate data gathered during different reward-related experiences, thus enabling researchers to capture greater variability of the data.

### 4.5. Conclusions and future directions

The present study set out to develop and validate a generic model of the ongoing fluctuations in the BOLD signal in the VS, using EEG alone. A series of analyses suggested that the developed EEG model, which relies on only eight scalp channels, was valid in predicting activation in the VS and related network across different sessions, individuals, and experimental conditions and was further modulated by reward value. Thus, this probe may be used in settings that preclude fMRI for basic-scientific as well as translational research for developing clinical applications, such as neurofeedback.

## Acknowledgments

We would like to thank all of the participants for their time. We also thank Kevin Larcher for his great support with the analysis of the pilot data, Shira Balter, Ilana Podlipski and Guy Gurevitch for their useful input regarding the computation of the model procedure.

## Funding

This work has received funding from the MOST-FRQNT-FRQS grant for collaborative research between Quebec and Israel on Biomedical Imaging. Additional funding was provided by a Foundation Grant to RJZ from the Canadian Institutes of Health Research, and from the Canada First Research Excellence Fund awarded to McGill University for the Healthy Brains for Healthy Lives initiative. RJZ is a fellow of the Canadian Institute for Advanced Research. This work was additionally supported by the European Union’s Horizon 2020 Framework Program for Research and Innovation under the Specific Grant Agreement No. 945539 (Human Brain Project SGA3) and by the Israel Science Foundation (grant No. 2923/20), within the Israel Precision Medicine Partnership program to TH. NS was supported by the Tel-Aviv University BrainBoost innovation center postdoctoral fellowship, and the Eldee Foundation and the Bloomfield family TAU-McGill collaboration scholarship. The funders had no role in the conceptualization, design, data collection, analysis, the decision to publish, interpretations, or preparation of the manuscript.

## Author contributions

NS, TH, AD, and RZ, conceptualized the research, and wrote the paper; NS, TB, TH, AD, and RZ designed the research; NS, ND, SN acquired the data, NS, TH, and GP designed the computation of the prediction model, GP implemented the computation prediction model, NS and MD analyzed the data.

## Notes

### Competing Interest Statement

TH is the Chief Medical Scientist in GrayMatters Health. TH, NS, GP, RZ, and AD have a pending patent (63/042,404). The authors declare no other competing financial or other interests.

