## Supplementary materials for "Development and validation of an fMRI-informed EEG model of reward-related ventral striatum activation"

**<u>Whole-brain VS-EFP correlates consistent across modeling and validation study cohorts</u>**

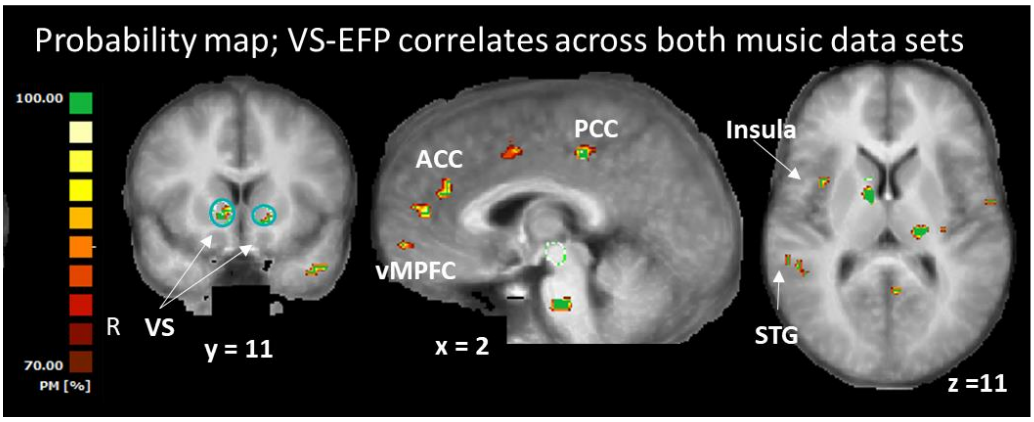

**Fig. S1. Whole-brain correlates of the VS-EFP that are consistent across musical-pleasure datasets.** The brain areas consistently correlated with the VS-EFP in both the modeling and validation study cohorts are depicted as a probability map constructed from the significant correlation across the two maps, each thresholded at the level resulting in ~80000 correlated voxels (see fig. 3; modeling: p<.001; validation: 0.00015).

### Neuroanatomical specificity of the EFP-model

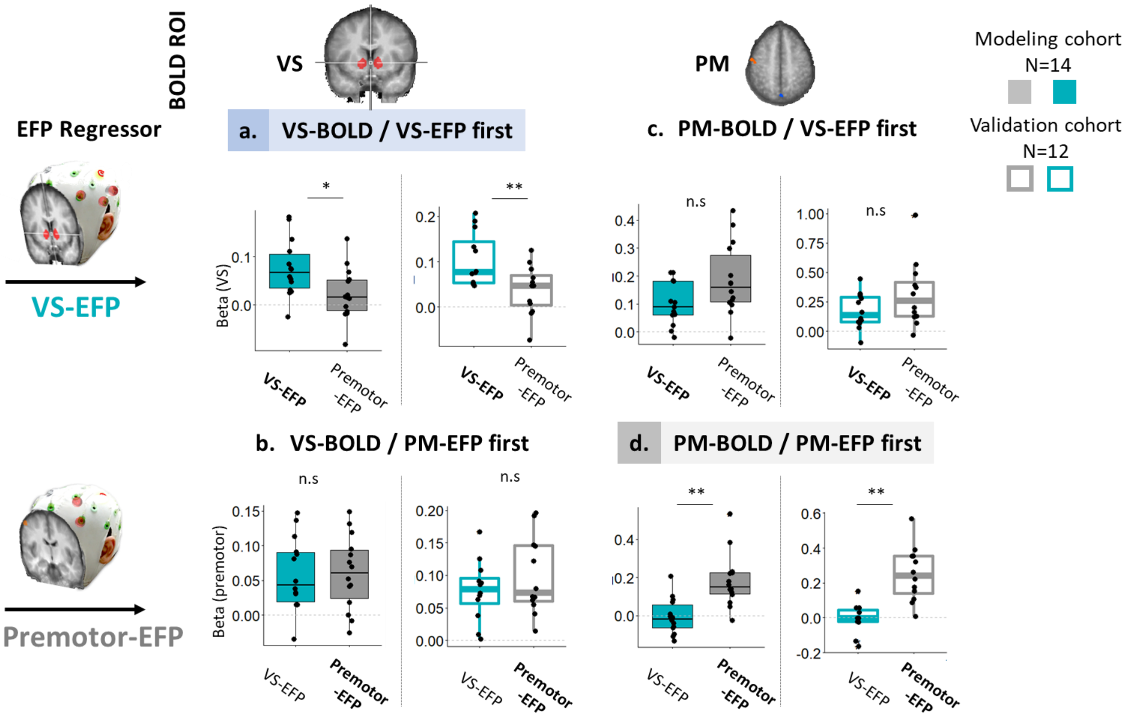

**Fig. S2. Neuroanatomical specificity of the EFP-model**. The neuroanatomical specificity of the VS-EFP (top panel) was assessed by comparing the correlates of the VS-EFP to the premotor-EFP (lower panel). Here, we depict the results of two GLM analyses applied for the two ROIs. Each GLM per ROI was applied while the first EFP, e.g., VS-EFP, was regressed with an orthogonalized version of the other EFP (e.g., premotor-EFP). As can be seen, the specificity of correlation between EFP and ROI is evident only for the congruent conditions (a.,d., ROI + EFP regressed first).

Abbreviations: VS = ventral striatum; PM = premotor; EFP; electrical fingerprint; BOLD: blood oxygenation level dependent

**Table S1. Whole brain correlates of VS-EFP.** Brain areas that were correlated with the VS-EFP consistently across both modeling and validation datasets, with a minimal cluster size of 125 voxels. The regions were extracted from a probability map of that was constructed from the whole-brain maps of the VS-EFP correlates of both datasets, and at a threshold of > 67% probability. Peak voxel within each cluster and the corresponding t-value are reported per dataset;

| Cluster size | Anatomical area | Dataset | Coordinates | | | t - value |
|---|---|---|---|---|---|---|
| | | | X | Y | Z | |
| 147 | R. MTG, BA 37/21c | Modeling | 57 | -49 | 1 | 6.879565 |
| | | Validation | 60 | -46 | 4 | 6.72157 |
| 151 | R. MTG , BA 21i | Modeling | 57 | -10 | -12 | 5.292744 |
| | | Validation | 57 | -7 | -13 | 7.612225 |
| 498 | R. posterior STG, BA 22c | Modeling | 46 | -43 | 10 | 4.825942 |
| | | Validation | 54 | -33 | 7 | 10.21706 |
| 237 | R. anterior STG, BA 22r | Modeling | 51 | 2 | -8 | 5.60867 |
| | | Validation | 51 | 2 | -8 | 13.38176 |
| 132 | R. OFC, BA12/47 | Modeling | 39 | 32 | 1 | 5.603713 |
| | | Validation | 39 | 35 | 0 | 7.97907 |
| 179 | R Insula, BA 13 | Modeling | 36 | 6 | 8 | 4.580179 |
| | | Validation | 33 | 8 | 10 | 7.705208 |
| 478 | R. amygdala (LB) | Modeling | 27 | 0 | -23 | 5.938266 |
| | | Validation | 30 | 2 | -20 | 8.506200 |
| 2117 | R. Cerebellum, anterior lobe | Modeling | 18 | -49 | -20 | 6.397427 |
| | | Validation | 27 | -34 | -20 | 9.839035 |
| 385 | R. Amygdala | Modeling | 30 | -4 | -14 | 4.860993 |
| | | Validation | 30 | -4 | -11 | 7.906753 |
| | R. Putamen | Modeling | 30 | -1 | -8 | 5.073108 |
| | | Validation | 33 | -1 | -2 | 6.6660178 |
| 149 | R. Cerebellum, Crus I | Modeling | 33 | -64 | -32 | 5.791658 |
| | | Validation | 26 | -70 | -32 | 6.109346 |
| 329 | R. Putamen | Modeling | 25 | -12 | -5 | 5.62418 |
| | | Validation | 27 | -10 | -5 | 9.637306 |
| 538 | R. VS | Modeling | 6 | 9 | 1 | 5.604451 |
| | | Validation | 9 | 8 | 1 | 7.710344 |
| 254 | R. Anterolateral Thal. | Modeling | 9 | -2 | 7 | 5.266632 |
| | | Validation | 9 | -1 | 10 | 7.824749 |
| 217 | R. Precuneus | Modeling | 12 | -43 | 46 | 5.685361 |
| | | Validation | 12 | -43 | 46 | 7.154764 |
| 580 | R. PCC, BA 31 | Modeling | 6 | -28 | 37 | 5.435513 |
| | | Validation | 0 | -32 | 40 | 7.042501 |
| 520 | R. pons | Modeling | 3 | -19 | -29 | 5.326486 |
| | | Validation | 3 | -19 | -32 | 8.819175 |
| 410 | vmPFC, BA10 | Modeling | 3 | 50 | 17 | 5.319791 |
| | | Validation | 3 | 43 | 16 | 7.575456 |
| 183 | R.+ L. vmPFC, BA11/10 | Modeling | 3 | 53 | -2 | 6.411053 |
| | | Validation | 3 | 56 | -2 | 7.230756 |
| 358 | R. ACC, BA 24 | Modeling | 1 | -1 | 39 | 5.190372 |
| | R. dorsal ACC, BA 32 | Validation | 0 | 6 | 49 | 6.488818 |
| 372 | R. dorsal ACC, BA32 | Modeling | 0 | 38 | 25 | 5.158216 |
| | | Validation | 0 | 35 | 25 | 7.38122 |
| 164 | L. dorsal PCC, BA31 | Modeling | -6 | -52 | 10 | 4.99206 |
| | | Validation | -6 | -52 | 13 | 8.109638 |

| | | | | | | |
|---|---|---|---|---|---|---|
| 363 | L. vmPFC, BA 32 | Modeling | -2 | 43 | 1 | 5.109111 |
| | | Validation | -6 | 44 | 1 | 7.757345 |
| 236 | L. PCC, BA31 | Modeling | -12 | -34 | 37 | 6.173559 |
| | | Validation | -9 | -34 | 38 | 7.466651 |
| 453 | L. Cerebellum, posterior lobe | Modeling | -6 | -58 | -32 | 6.300318 |
| | | Validation | -6 | -64 | -35 | 7.503042 |
| 856 | L. VS | Modeling | -12 | 11 | -2 | 4.573865 |
| | | Validation | -15 | 8 | -2 | 9.029176 |
| | L. anterior Insula, BA 13 | Modeling | -30 | 26 | 1 | 6.256996 |
| | | Validation | -33 | 26 | 1 | 6.681299 |
| | L. GPi | Modeling | -12 | -7 | -5 | 5.903521 |
| | | Validation | -15 | -7 | -2 | 8.182054 |
| 436 | L. Claustrum | Modeling | -24 | -22 | 16 | 5.634065 |
| | | Validation | -24 | -19 | 19 | 7.348473 |
| | L. Thal., posterior lateral | Modeling | -22 | -19 | 10 | 5.373977 |
| 1074 | L. Amygdala | Modeling | -24 | -10 | -5 | 7.082157 |
| | | Validation | -27 | -8 | -12 | 8.287025 |
| | L. Putamen | Modeling | -30 | -19 | 7 | 5.573330 |
| | | Validation | -30 | -16 | 1 | 7.279988 |
| 185 | L. Cerebellum, anterior lobe | Modeling | -18 | -55 | -20 | 5.037019 |
| | | Validation | -22 | -56 | -20 | 6.61462 |
| 418 | L. Cerebellum, anterior lobe | Modeling | -29 | -40 | -26 | 5.663418 |
| | | Validation | -27 | -43 | -23 | 7.3476 |
| 497 | L. Fusiform Gyrus, BA 37/19 | Modeling | -39 | -64 | -12 | 5.841224 |
| | | Validation | -42 | -64 | -5 | 8.090832 |
| 481 | L. Temporal pole, BA 38 | Modeling | -36 | 11 | -29 | 5.210886 |
| | | Validation | -42 | 8 | -25 | 9.297838 |
| 242 | L. Superior Temporal Gyrus, BA 22 | Modeling | -45 | -43 | 23 | 5.446657 |
| | | Validation | -48 | -43 | 19 | 6.512595 |
| 538 | L. Middle Temporal Gyrus, BA 21 | Modeling | -54 | -25 | -2 | 7.44871 |
| | | Validation | -54 | -22 | 1 | 7.665364 |
| 129 | Superior Temporal Gyrus, BA 21 | Modeling | -49 | 2 | -11 | 4.695522 |
| | | Validation | -48 | 2 | -11 | 7.086081 |
| 160 | L. Sub-central Area, BA 43 | Modeling | -58 | -5 | 10 | 5.129484 |
| | | Validation | -54 | -1 | 7 | 7.220856 |

Talairach coordinates are used. Cluster size denotes number of voxels per region. In cases that the cluster included more than one anatomical area, several peak voxels are reported. Brain regions that fall within grey matter are reported. **Abbreviations**: BA = Brodmann Area; MTG = Middle Temporal Gyrus; STG = Superior Temporal Gyrus; OFC = OrbitoFrontal Cortex; VS = Vental Striatum; Thal. = Thalamus; PCC = Posterior Cinculate Cortex; vmPFC = ventromedial PreFrontal Cortex; ACC = Anterior Cingulate Cortex. GPi = Globus Pallidus internal

**Table S2. Whole brain VS-EFP neuroanatomical specificity**. Brain areas that were correlated with the VS-EFP, more than with premotor EFP consistently across both modeling and validation study datasets, with a minimal cluster size of 125 voxels. The regions were extracted from a probability map of that was constructed from the whole-brain maps of the contrast VS-EFP > premotor-EFP of both datasets, and at a threshold of > 67% probability. Peak voxel within each cluster and the corresponding t-value are reported per dataset;

| Cluster size | Anatomical area | Coordinates | | | t - value |
|---|---|---|---|---|---|
| | | X | Y | Z | |
| 484 | R. ACC, BA32 | 0 | 33 | 19 | 4.755487 |
| 484 | R. ACC, BA32 | 6 | 29 | 22 | 8.549871 |
| 239 | R. DS, caudate body | 9 | -1 | 16 | 4.238058 |
| 239 | R. DS, caudate body | 12 | 4 | 13 | 7.022169 |
| 162 | R. VS | 12 | 11 | 1 | 3.516539 |
| 162 | R. VS | 9 | 10 | 1 | 6.003746 |

Talairach coordinates are used. Cluster size denotes number of voxels per region. Brain regions that fall within grey matter are reported. **Abbreviations**: BA = Brodmann Area; DS = Dorsal Striatum; VS = Vental Striatum; ACC = Anterior Cingulate Cortex.

**Table S3. Whole brain correlates of VS-EFP during the MID task.** Brain areas that were correlated with the VS-EFP during the MID task. Brain regions correlating with the VS-EFP signal, thresholded at FDR-corrected *p* <.005 with a threshold of 125 cluster size.

| Cluster size | Anatomical area | Coordinates | | | t - value |
|---|---|---|---|---|---|
| | | X | Y | Z | |
| 2854 | R. Precentral Gyrus, BA4, M1 | 57 | -13 | 31 | 6.453201 |
| 405 | R. anterior Insula, BA 13 | 48 | 11 | 1 | 5.700069 |
| 199 | R. MTG, BA 37 | 45 | -64 | 7 | 5.366115 |
| 1223 | R. Insula, BA 13 | 30 | -7 | 16 | 7.084366 |
| 804 | R. anterior Insula, BA 13 | 39 | 26 | 7 | 6.254556 |
| 244 | R. Precentral Gyrus, BA 6, premotor cortex | 30 | -13 | 61 | 5.246312 |
| 244 | R. posterior Insula, BA 13 | 33 | -31 | 19 | 6.706285 |
| | R. Cerebellum, anterior lobe | 12 | -55 | -17 | 8.492515 |
| | R. Cuneus, BA 17 | 6 | -76 | 10 | 8.414027 |
| | R. Pons | 0 | -22 | -26 | 8.358302 |
| | L. Pons | -6 | -19 | -29 | 7.269628 |
| | L. Cerebellum, anterior lobe | -9 | -67 | -8 | 9.084085 |
| | L. Cuneus, BA 18 | -21 | -82 | 22 | 7.21393 |
| 52464 | L. Lingual Gyrus, BA 17 | -18 | -88 | 1 | 7.41414 |
| | R. Putamen | 27 | -7 | -2 | 7.98452 |
| 3016 | L. midbrain, VTA | -6 | -16 | -8 | 7.722530 |
| 199 | R Thal., Pulvinar | 21 | -31 | 10 | 6.375166 |
| 565 | Post central gyrus, BA 3, somatosensory | 21 | -31 | 64 | 6.094884 |
| | L. Caudate body | -21 | -19 | 22 | 7.624059 |
| | R. Thal. | 6 | -13 | 16 | 6.826271 |
| | L. Thal. | -15 | -16 | 16 | 7.200933 |
| | L. Putamen | -27 | -4 | -8 | 7.483016 |
| | L. Insula, BA 13 | -33 | -10 | 13 | 7.070439 |
| | L. transverse temporal gyrus, BA 41/42 | -54 | -19 | 13 | 7.549177 |
| 20746 | L. Inferior Parietal Lobule, BA 40 | -54 | -40 | 43 | 6.871533 |
| 500 | R. ACC, BA32 | -3 | 17 | 34 | 5.709224 |
| 315 | L. dorsal PCC, BA 31 | -9 | -22 | 43 | 6.637782 |
| 164 | L. VS | -9 | 14 | 7 | 5.45807 |
| 3888 | L. Precuneus, BA40 | -24 | -43 | 49 | 7.547597 |
| 187 | L. Cerebellum | -15 | -64 | -36 | 5.410328 |
| 1112 | L. anterior Insula, BA 13 | -33 | 26 | 7 | 6.695986 |
| 176 | L. Superior parietal lobule, BA7 | -27 | -58 | 43 | 6.929028 |
| 188 | L. MFG, BA 10 | -30 | 47 | 16 | 5.489028 |
| 173 | L. STG, BA 22 | -51 | 8 | -2 | 5.699324 |

Talairach coordinates are used. Cluster size denotes number of voxels per region. In cases that the cluster included more than one anatomical area, several peak voxels are reported. Brain regions that fall within grey matter are reported. **Abbreviations**: BA = Brodmann Area; VTA = Ventral Tegmental Area, MTG = Middle Temporal Gyrus; VS = Vental Striatum; Thal. = Thalamus; PCC = Posterior Cinculate Cortex; ACC = Anterior Cingulate Cortex.
